# Needle in a haystack: culturing plant-beneficial Helotiales lineages from plant roots

**DOI:** 10.1101/2024.11.15.623725

**Authors:** Pauline Bruyant, Jeanne Doré, Laurent Vallon, Yvan Moënne-Loccoz, Juliana Almario

## Abstract

Root-associated Helotiales fungi are increasingly recognized as beneficial fungal partners promoting plant growth under nutrient-limited conditions, particularly in non-mycorrhizal hosts, lacking the ancestral arbuscular mycorrhizal symbiosis. However, the ecology of these fungi is still cryptic as relatively few lineages have been successfully cultivated from roots for further study. Here, we attempted the mass isolation of root endophytic fungi to evaluate the recovery of known plant-beneficial Helotiales lineages using a tailored culture-based approach. We sampled six wild non-mycorrhizal species from the Brassicaceae, Caryophyllaceae, and Cyperaceae, growing in nutrient-limited alpine soils. We isolated 602 root endophytes and compared this culturable diversity with the one observed via fungal ITS2 metabarcoding. Metabarcoding revealed that Helotiales taxa dominated the fungal communities, with 43% of these detected taxa also represented in our collection. Accordingly, most root endophytes in our collection (53%) were Helotiales. These isolates, some with P solubilisation potential, belonged primarily to three Helotialean clades and were phylogenetically related to plant growth-promoting or mycorrhizal-like strains. This analysis highlights that roots of alpine non-mycorrhizal plants are reservoirs of plant-beneficial root-endophytic Helotiales, and the isolates obtained are a promising resource to explore the plant-beneficial mechanisms and ecological traits of these fungi.

## Introduction

The microbiota can mediate host’s adaptation to harsh environmental conditions (Petipas *et al*., 2021). In the case of plants, root-associated microorganisms have been found to promote plant growth under unfavourable environmental conditions such as drought, high salinity and low nutrient levels (Almario *et al*., 2017; Fitzpatrick *et al*., 2019; Liu *et al*., 2022). Exploring the root microbiota of plants growing in nutrient-limited soils has led to the description of new plant-beneficial microbial lineages, including mycorrhizal-like fungi capable of transferring nutrients like N and P to their host without forming specific symbiotic structures in plant roots (Almario *et al*., 2022; Bruyant *et al*., 2024).

Most plants rely on the arbuscular mycorrhizal symbiosis to meet their phosphorus (P) requirements (Brundrett and Tedersoo, 2018), but several plant lineages have abandoned this symbiosis (Werner *et al*., 2018). Prominent non-mycorrhizal (Non-AM) families include the Brassicaceae, Caryophyllaceae, and Cyperaceae (Werner *et al*., 2018). Until recently, no alternative nutritional strategy was described for these plants, but recent research (Almario *et al*., 2022; Bruyant *et al*., 2024) suggests that some of these Non-AM plants may depend instead on new associations with mycorrhizal-like root-endophytic fungi from the Helotiales order.

Root-associated Helotiales include plant-beneficial lineages that can improve plant nutrition as symbiotic partners in ericoid mycorrhizae or ectomycorrhizae, and as non-symbiotic beneficial root endophytes, including mycorrhizal-like fungi capable of nutrient transfer (Grelet *et al*., 2009; Tedersoo and Smith, 2013; Almario *et al*., 2017; Kowal *et al*., 2018; Hill *et al*., 2019). However, for most root-associated taxa their ecology remains cryptic (Wang *et al*., 2006; Quandt and Haelewaters, 2021; Bruyant *et al*., 2024). Mycorrhizal-like Helotiales belong to two phylogenetically-distant clades, which suggests that the capacity to transfer nutrients to plants is widespread within this order (Bruyant *et al*., 2024). These fungi are particularly prevalent in plants from nutrient-poor soils (Toju *et al*., 2016; Almario *et al*., 2017; Jumpponen *et al*., 2017; Botnen *et al*., 2020), such as those found in arctic and alpine environments (Weintraub, 2011; Vincent *et al*., 2014), suggesting they play a key role in mediating plant adaptation by supporting plant nutrition under these extreme conditions. These associations could be even more critical for Non-AM plants growing in these environments and unable to rely on the ancestral AM symbiosis to improve their nutrition. Other plant-beneficial Helotiales, such as those from the *Phialocephala* clade, as well as *Cadophora* and *Lachnum* strains, have been shown to enhance biomass and shoot nutrient content in various plant species (Jumpponen *et al*., 1998; Bizabani and Dames, 2015; Surono and Narisawa, 2017, 2021), via the mineralization of organic N and P sources (Della Monica *et al*., 2015; Surono and Narisawa, 2021; Mikheev *et al*., 2022), solubilisation of N or P (Mikheev *et al*., 2022) or unknown mechanisms (Vohník *et al*., 2005). However, the ecology of plant-beneficial Helotiales, and especially of mycorrhizal-like lineages, remains poorly understood. Key aspects such as their prevalence, diversity, functions and genes involved in plant nutrition are still largely unknown. To address these issues, it is essential to isolate and study a wide range of fungal strains, aiming to capture the full diversity of these associations. Only a few isolates have been obtained so far, and we hypothesized that culture conditions can be designed to recover a much broader range of these fungi.

The aim of this study was to characterize the taxonomic diversity of culturable Helotiales and especially of suspected plant-beneficial Helotiales lineages, amongst root endophytes associated with alpine Non-AM plants. For this purpose, we sampled plants from wild-growing populations of two species of the Non-AM families Brassicaceae (i.e., *Arabis alpina* and *Draba azoides*), Cyperaceae (*Carex sempervirens* and *Carex curvula*) and Caryophyllaceae (*Minuartia verna* and *Silene acaulis*) in two sites from the French Alps characterized by low nutrient availability. Fungi were isolated from roots and characterized by ITS sequencing, and findings were compared with ITS metabarcoding data. In addition, we assessed the phylogeny of culturable Helotiales fungi given their prevalence in metabarcoding analyses. Finally, phosphate solubilisation potential was assessed *in vitro* for selected isolates, with the hypothesis that this capacity might be prevalent in root Helotiales considering their potential to enhance P nutrition of Non-AM plants.

## Experimental procedures

### Plant sampling in natural conditions

To characterize the fungal root microbiota six individuals from each of the three plant species *Arabis alpina* (Brassicaceae), *Carex sempervirens* (Cyperaceae), and *Minuartia verna* (Caryophyllaceae) were sampled from the Col du Galibier, and six individuals from each of *Draba azoides* (Brassicaceae), *Carex curvula* (Cyperaceae), and *Silene acaulis* (Caryophyllaceae) were collected from the Col du Lautaret in June and July 2020 within a 250 m^2^ area at each site. To isolate fungi from plant roots three individuals from each of these plant species were obtained from the same populations during July 2021 or July 2022. The two sites are located only 2 km apart in the Alps (Auvergne Rhône-Alpes region, France) but have a different geological origin and soil physico-chemistry (Table S1). Entire plants were sampled, roots were separated from the surrounding soil and thoroughly washed with ultra-pure water. Cleaned roots were then used for isolation of fungi (plants sampled in 2021 and 2022) or DNA extraction followed by ITS metabarcoding (plants sampled in 2020) as described below.

### Isolation of fungi from surface-disinfected roots of Non-AM plants

After washing, roots were surface disinfected in ethanol 70% for 90 s, then in bleach 3% for 4 min, and rinsed in sterilized ultrapure water three times. Therefore, it is likely that most root fungi obtained corresponded to true root endophytes. Equivalent numbers of root pieces (0.5 cm each; 3,920 root fragments in total) were plated on (1) 0.5× strength Potato Dextrose Agar (PDA-50), (2) Hagem’s agar (HgM; Walker *et al*., 2011), or (3) Minimal Medium (MM (Almario *et al*., 2017. PDA is a rich medium, MM a minimal medium, and HgM targets specifically the slow-growing fungi (Walker *et al*., 2011). In July 2021, 720 root fragments were plated on media MM or PDA-50, whereas in July 2022 3,200 fragments were plated on MM, PDA-50 or HgM (Table S2). Since plant species displayed contrasted sizes, they differed in root length available for sampling, and the number of 0.5-cm root fragments that could be used for plating varied from 488 for the Brassicaceae *D. azoides* to 1262 for the Cyperaceae *C. curvula* (Table S2). Half the plates were kept at room temperature and the others at 4°C to select for cold-growing fungi, for up to three months. Dates of appearance and visual properties of fungal colonies were recorded. We discarded any yeast-like colonies as we were interested in filamentous fungi, as well as the few (<5%) *Penicillium*-like and *Aspergillus*-like fungi (based on typical colony morphology) as they are very often lab contaminants. We isolated representative colonies of every morphology in each plant species × medium combination, purified them by three successive cultures on PDA (all grew on this medium), and then characterized the purified isolates by sequencing the ITS2 region of the rDNA (see below).

### Storage of the fungal collection

All sequenced isolates were stored on PDA plates at 4°C and as PDA plugs in 50% glycerol at -80°C. A comparative test involving 30 different Helotiales isolates indicated that the glycerol storage method at -80°C was superior to the one at 4°C for long-term preservation. Indeed, all tested Helotiales could be revived without contamination after one year of storage at -80°C, whereas fungi stored on PDA plates at 4°C failed to grow after this period.

### ITS2 sequencing and isolate identification

To extract DNA, a 0.5 cm^3^ PDA plug of each isolate was put in an autoclaved Fastprep tube (MP Biomedicals, Irvine, CA, USA) whose conical base was filled with 106-μm glass beads (Sigma-Aldrich, Darmstadt, Germany). Lysis buffer (400 mM Tris-HCl pH 8, 60 mM EDTA pH 8, 150 mM NaCl, 1% SDS) was added. Plugs were ground using a FastPrep-24 5G (MP Biomedicals) for 1 min at 6 m/s and incubated for 5 min at room temperature, followed by an additional round of grinding. 150 μL of 3M potassium acetate was added, tubes were centrifuged to eliminate cell debris, and supernatants were transferred to new tubes containing one volume of isopropanol. The tubes were inversed to facilitate mixing, and centrifuged. Supernatants were removed and DNA-containing pellets were washed with 70% ethanol. The tubes were centrifuged, supernatants were discarded, and the tubes were let to dry for 10 min at room temperature before resuspending pellets in sterile ultrapure water. DNA was then used for ITS2 amplification using primers fITS7 (Ihrmark *et al*., 2012) and ITS4 (Gardes and Bruns, 1993) at 0.8 μM each. Reaction mix was made with the Hot FirePol blend master mix with 12.5 mM MgCl2 (5×) (Solis BioDyne, Tartu, Estonia). Genomic DNA was diluted at 1:10 and 1 μL of diluted DNA was used. PCR cycling conditions were 95°C for 3 min followed by 35 amplification cycles (95°C for 30 s, 55°C for 30 s, 72°C for 45 s), followed by a final elongation at 72°C during 5 min. When PCR failed, the full ITS region was amplified using the primers ITS4-ITS1 (White *et al*., 1990) or ITS4-ITS5 (Bertini *et al*., 1999), following the same PCR steps but with an annealing temperature at 54°C. PCR products were verified by electrophoresis before Sanger sequencing (Microsynth AG, Vaulx-en-Velin, France). When indicated, primers ITS4 and ITS5 were used to amplify the complete ITS region for 30 of 319 Helotiales isolates selected to represent the diversity observed from ITS2 sequencing on all the 319 Helotiales isolates obtained. ITS sequences were taxonomically classified using the UNITE fungal database (UNITE version of 29/11/2022) (Nilsson *et al*., 2019; Abarenkov *et al*., 2022) in Mothur v.1.44.3 (Schloss *et al*., 2009). ITS2 sequenced were also clustered in OTUs using cluster.seqs with dgc method, with a 0.03 threshold in Mothur v.1.44.3 (Schloss *et al*., 2009). Sanger sequencing data was submitted to NCBI GenBank (see Table S1).

### ITS comparison of isolates and metabarcodes

For fungal ITS2 metabarcoding, root DNA was extracted and used for ITS2 PCR amplification using primers fITS7 and ITS4, before sequencing in an Illumina Miseq (2 × 300 b), as described in Almario *et al*. (2017). ITS2 amplicon sequences were processed and analysed using ITSx (Bengtsson-Palme *et al*., 2013) and Mothur as described in Almario *et al*. (2017). ITS2 metabarcoding data is available under project PRJNA1141433 Biosamples SAMN42888810 to SAMN42888827 and SAMN42888846 to SAMN42888863 in the NCBI database.

Sanger ITS2 sequences from isolates were then compared with ITS2 amplicon sequences obtained from the same plant populations (but not the same individuals). ITSx (Bengtsson-Palme *et al*., 2013) was also to extract the ITS2 region from all Sanger sequences from the isolates. Then, they were aligned against the representative OTU metabarcoding sequences (from all OTUs appearing in root samples defined at 99% threshold) using MAFFT (method FFT-NS-2 and default parameters, including the option to leave gappy regions (Katoh *et al*., 2002; Katoh and Standley, 2013)). A metabarcoding OTU was considered covered if at least one isolate matched the sequence with more than 97% identity. An isolate could cover several metabarcoding OTUs as they were defined at 99% identity threshold. ITS2 and full size ITS sequences from the isolates are available under GenBank accession numbers PQ323580 to PQ324178 and PQ286942 to PQ286971 in the NCBI database (see table S2).

### Phylogenetic tree of Helotiales isolates

To further elucidate the phylogenetic diversity of root-endophytic Helotiales isolated from alpine Non-AM plants, 30 of 319 Helotiales isolates were selected to represent the overall diversity observed among isolates (based on ITS2 sequencing). The 30 isolates harboured ITS sequences sharing less than 92% sequence similarity. Complete ITS sequences from these 30 isolates were aligned with those from reference strains, from strains with plant growth-promoting or nutrient acquisition-promoting capacities, and from lichen-associated strains, as described in Bruyant *et al*. (2024). For each of the 30 isolates, the relative abundance of their ITS variants, in all sequences of the collection and in the corresponding sequences from metabarcoding analysis, was determined based on 97% similarity in the ITS2 sequence.

### Testing solubilisation of P *in vitro*

To assess the P solubilization potential of a diverse range of fungi, 142 isolates (listed in Table S2) derived on MM or PDA-50 from the roots of six plant species sampled in 2021 were used. They were tested using Pikovskaya’s (PVK) medium, which contains calcium phosphate (5 g/L) (Pikovskaia, 1948). Each of the 142 isolates was inoculated by placing 0.5 mm^3^ of culture onto PVK agar plates (in triplicate). The plates were incubated at room temperature for up to 14 days. Phosphorus solubilization was determined by the presence of clear zones around the colonies, indicative of the conversion of insoluble calcium phosphate into soluble forms.

## Results

### Sampling effort and taxonomic diversity of root-endophytic fungi from alpine Non-AM plants

Only 1,733 of the 3,920 surface-sterilized root fragments (i.e. 44%) gave fungal colonies on plates (40% were obtained on MM, 31% on PDA-50 and 29% on HgM). The 1,733 isolates obtained were grouped based on colony morphology and time of colony formation, and 602 representatives of each group for each medium were kept for ITS2 sequencing. Half of the 1,733 isolates (51%) and of the 602 ones used for sequencing (47%) came from the Cyperaceae *C. sempervirens* and *C. curvula*, which was expected due to the higher availability of root material from these two species (Table S3).

At phylum level, the 602 isolates included 595 Ascomycota (98.8%), 3 Basidiomycota, 2 Mortierellomycota and 2 unclassified fungi. They corresponded to 19 orders, including a majority of Helotiales (53%), Pleosporales (25%) and Hypocreales (10%) (Fig. 1A). At the species level, 270 isolates were identified, with a confidence level exceeding 80%. The rarefaction curves indicated that a plateau was not reached, meaning that the entire diversity had not been recovered (Fig. 1B). Based on an extrapolation to twice the number of isolates, a plateau would not have been reached still, for each plant species (Inext package extrapolation function). Different levels of fungal diversity were recovered in the three plant species (number of OTUs at 97% sequence similarity; Chi^2^ tests, *P* = 0.014), with both Cyperaceae species showing higher levels of richness (Fig. 1B). However, contrary to expectations, similar levels of fungal diversity were recovered on the three media when considering each plant species independently (Chi^2^ tests on OTU numbers, *P >* 0.05).

**Figure 1.**
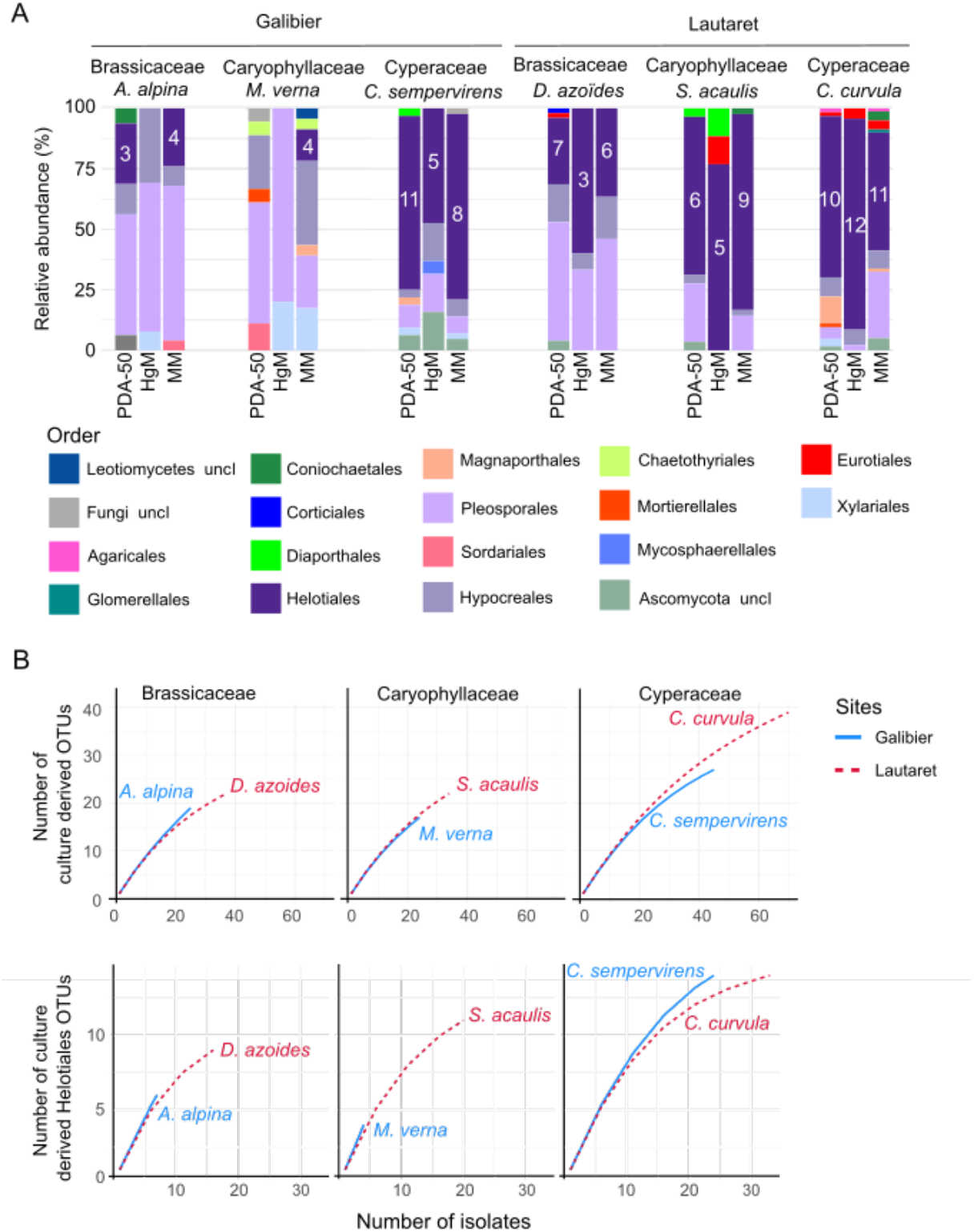
Taxonomic coverage of the fungal collection. (A) Relative abundance of each fungal order in the collection for each plant species and laboratory medium. White numbers indicate the number of different cultured Helotiales OTUs (clustered at 97% similarity) for each species and medium. Abbreviations: PDA-50, half-strength Potato Dextrose Agar; HgM, Hagem agar; MM, Minimal Medium; uncl, unclassified. (B) Rarefaction curves for each plant species based on all (top) or only Helotiales (bottom) culture-derived OTUs.

When investigating the diversity among the 319 isolated Helotiales, we documented 27 different OTUs (at 97% sequence similarity), which surpassed the diversity observed in other orders, including highly prevalent Pleosporales (23 OTUs), but a rarefaction plateau was still not reached (Fig. 1B). Similar Helotiales diversity was recovered in all plants (similar numbers of OTUs, chi^2^ tests, *P* > 0.05; Fig. 1A), but Helotiales were more frequently isolated from *C. curvula, C. sempervirens* (Cyperaceae) and *S. acaulis* (Caryophyllaceae) (Chi^2^ tests, *P* < 0.05). Moreover, when considering each plant species individually, we could see that only MM was effective at recovering Helotiales from all plant species sampled, as none were obtained with HgM and PDA-50 for *M. verna* and none with HgM for *A. alpina* (Fig. 1A).

In summary, mass isolation of root endophytes from alpine Non-AM plants revealed a diverse fungal community predominantly composed of Ascomycota, with a notable representation from the Helotiales order. MM emerged as the only medium effective for recovering Helotiales from all conditions tested.

### Comparison of the culture collection with metabarcoding OTUs

Based on the ratio of cultured fungal isolate numbers to metabarcoding OTUs, it is estimated that around 87% of root-associated taxa cannot be cultured (Wu *et al*., 2019). To estimate how well our isolate collection represented real life root-associated fungal communities, we compared our collection to the fungal diversity observed through ITS2 metabarcoding using a 97% sequence similarity threshold. From the 3277 metabarcoding fungal OTUs detected in root samples, 620 were present in our fungal collection (i.e. 18.9%). However, when considering the most abundant metabarcoding OTUs, i.e. 197 OTUs with top relative abundances representing 80% of the root fungal community, as much as one third of them was recovered as isolates in our collection (61 of 197, i.e. 31.1%). In turn, these taxa recovered represented approximately one third (35.0%) of the overall root fungal community, with as many as 7 of the 10 most abundant OTUs (including 6 Helotiales) represented by at least one isolate (Fig. 2).

**Figure 2.**
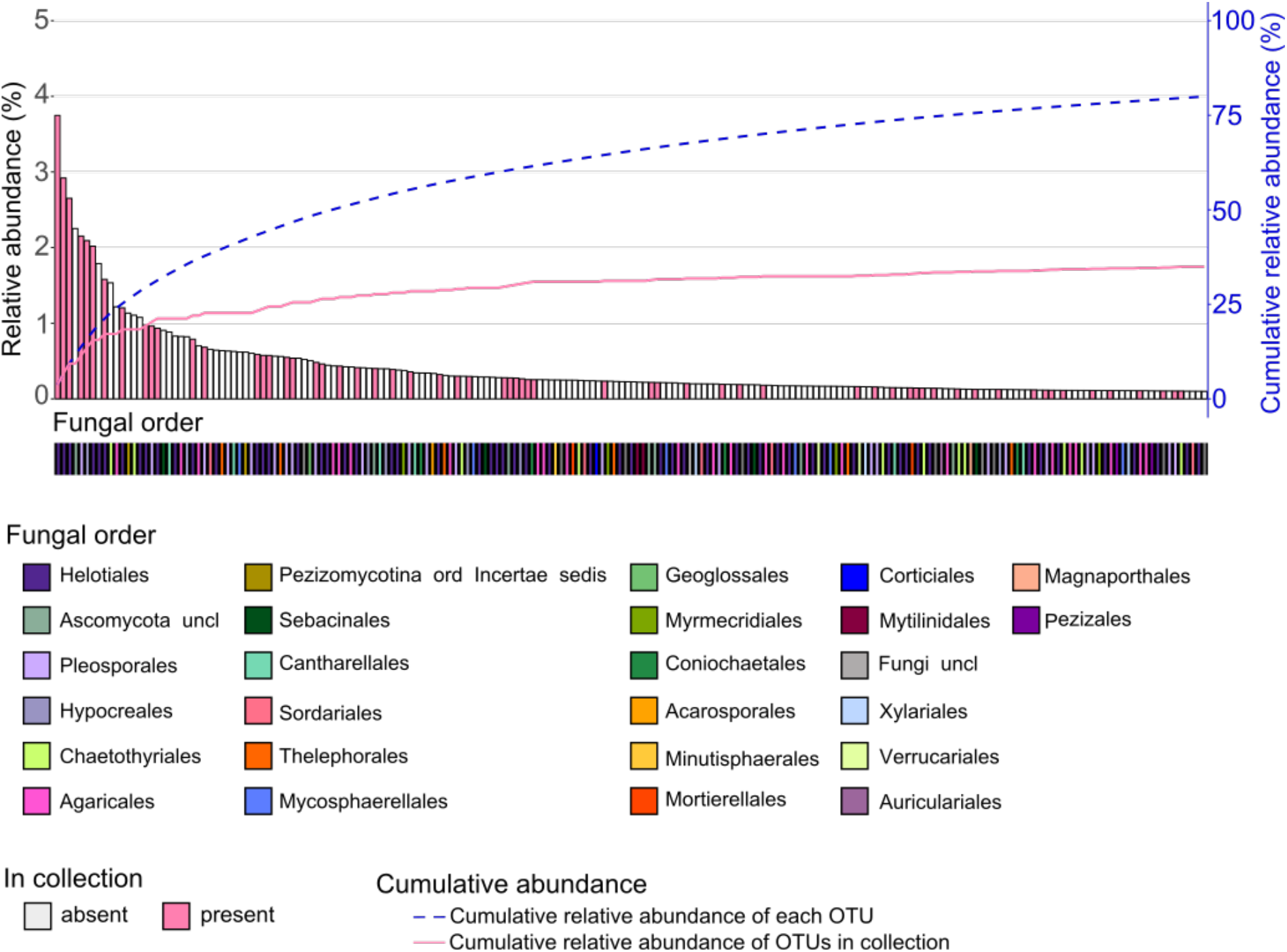
Representativeness of the culture collection by comparison with metabarcoding OTUs. The rank abundance plot shows the 197 most abundant root-associated fungal OTUs of Non-AM plants growing at Galibier and Lautaret together with their cumulative relative abundance (blue dashed line), representing a total of 80% of the relative abundance of all OTUs. For visual clarity, the 3,081 other OTUs (of relative abundance < 20%) are not shown. OTUs are color-coded according to the presence of at least one representative isolate in the culture collection (at 97% ITS2 sequence similarity). The percentages of naturally-occurring OTUs recovered as pure cultures are indicated by the pink line. The taxonomy of each OTU (order) is indicated below the barplot.

Certain taxa were slightly overrepresented in the collection, such as the Eurotiales (1.6% vs 0.8% by metabarcoding), Xylariales (1.7% vs 0.5%) and Magnaporthales (1.7% vs 0.2%), whereas others were underrepresented (Basidiomycota: 1.2% vs 18.2% by metabarcoding) or even absent (Verrucariales or Mycosphaerellales, amounting to respectively 0.59% and 2.2% of the metabarcoding data). Additionally, certain taxa of the isolate collection (39 of 602, i.e. 6.0%) were not evidenced in the metabarcoding dataset (Fig. 3). They were either unclassified Ascomycota or belonged to the Diaporthales, Pleosporales, Hypocreales, Xylariales, Eurotiales, Corticiales or Helotiales orders. Apart from *Penicillium* and *Alternaria*, these taxa do not appear on the list of common lab contaminants but could be soil contaminants, as they belong to saprotrophic or phytopathogenic taxa. Alternatively, they could be root endophytes not detected in metabarcoding analysis due to their low abundance in the roots or sequencing biases.

**Figure 3.**
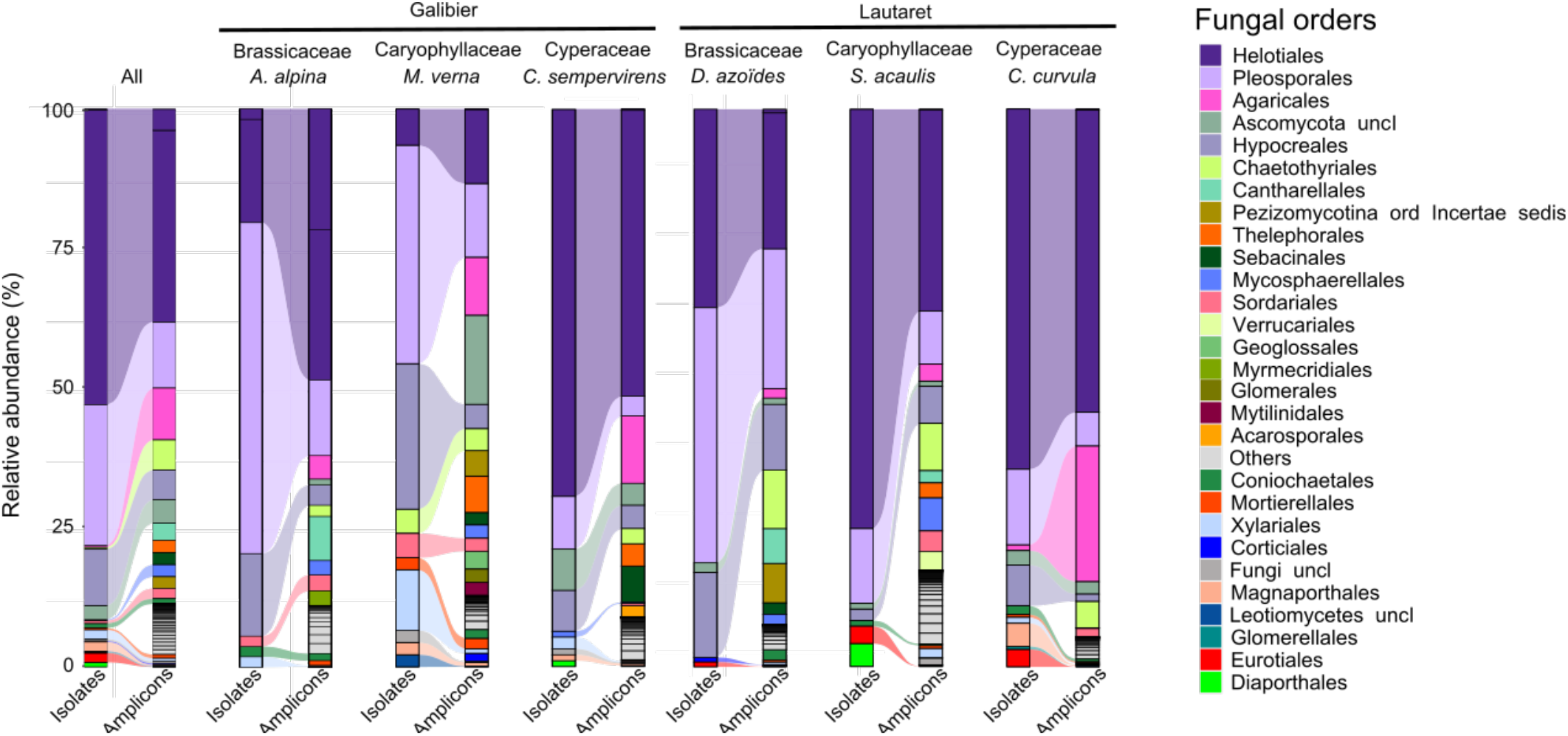
**Taxonomic diversity of the fungal root collection**, based on the comparison of fungal taxonomic composition at the order level between culture-dependent and culture-independent methods, for all plant species together (All) and each plant species separately. Isolates: taxonomy of the 602 fungi isolated from plant roots. Amplicons: taxonomy of fungal root-associated OTUs at the order level for each of the six plant species used to make the collection.

When focusing on the 1014 Helotiales OTUs found by metabarcoding (30.9% of the metabarcoding fungal OTUs), 434 of them (i.e. 42.8% of the Helotiales OTUs) were recovered in our isolate collection. This recovery rate is twice as high as the percentage recovered when considering the entire fungal community (i.e. 18.9%) and reached 55.9% when considering the most abundant taxa (top relative abundances representing 80% of the root fungal community) (Fig. 2). In comparison, for Pleosporales, the second most abundant order in both the collection and metabarcoding data, we recovered 93 of the total 281 metabarcoding OTUs (33.1% of the Pleosporales OTUs). Indeed, Helotiales were significantly overrepresented in our collection, comprising 53.0% of the isolates, compared with 30.9% of the OTUs in the metabarcoding data (or 38.2% of the total relative abundance).

In summary, our proposed culture method allowed the creation of a fungal collection with good representation of the most abundant fungi associated with the roots of Non-AM plants. The better coverage of Helotiales compared with other taxa shows that our cultivation process is particularly adapted for recovering diverse root-endophytic Helotiales.

### Phylogeny of root-endophytic Helotiales recovered from alpine Non-AM plants

Given the prevalence of the Helotiales order in our collection and its potential significance for plant adaptation to alpine environments, we examined the phylogeny of 30 Helotiales isolates, which represent the culture-based diversity identified in this study. These isolates fell into four of the six Helotiales clades: Hyaloscyphoid, Mollisioid, Pezizzeloid, and Discinelloid (Fig. 4).

**Figure 4.**
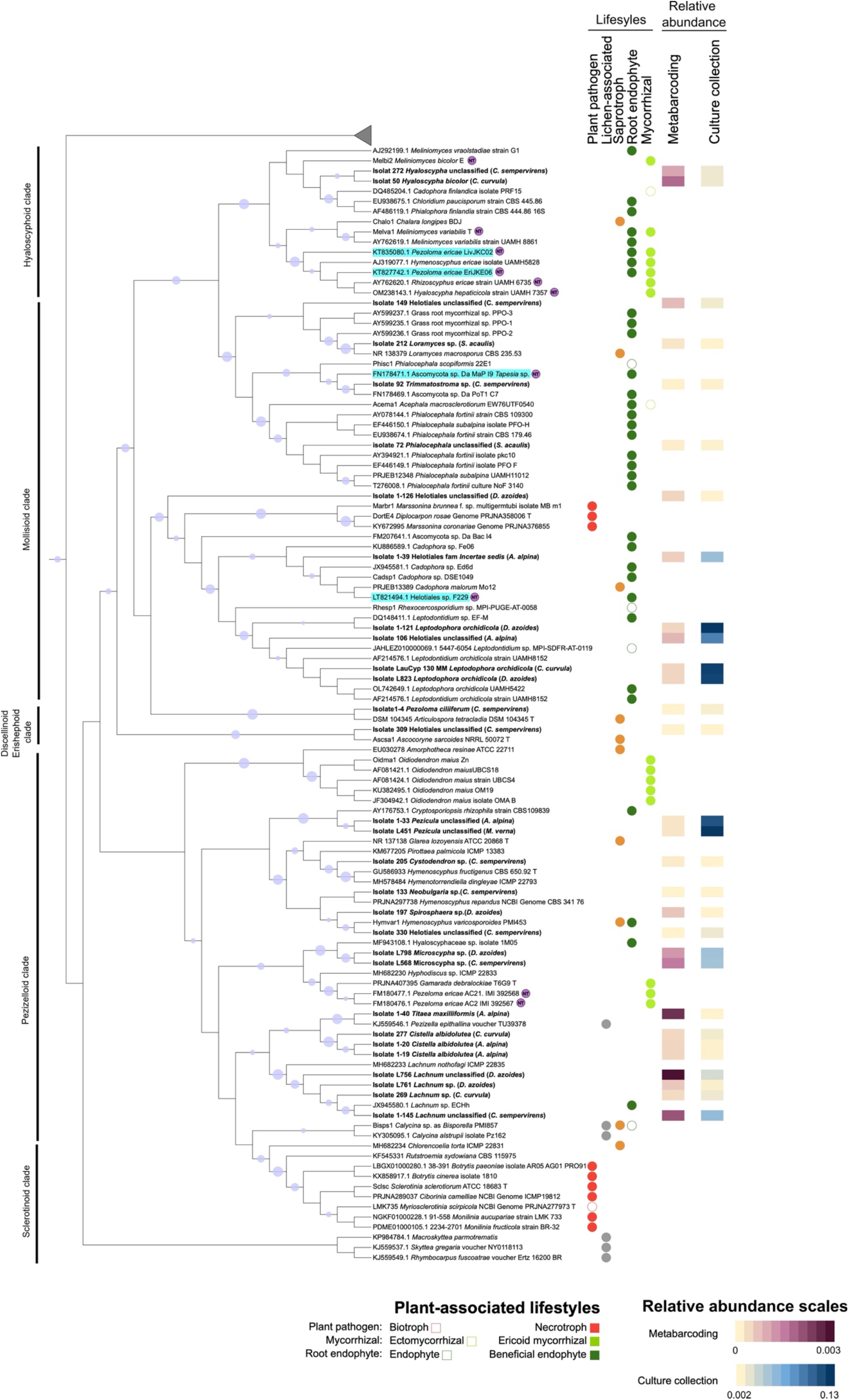
ITS2-based phylogenetic tree of representative Helotiales isolates (30, in bold) from Non-AM plants. ITS sequences from diverse Helotiales including reference strains, strains with plant growth-promoting or nutrient acquisition-promoting capacities, and from lichen-associated strains were included in the analysis described in Bruyant *et al*. 2024. “T” indicates a type strain. Helotiales phylogenetic clades described by Johnston *et al*. (2019) are indicated. The tree was inferred as described in Bruyant *et al*. 2024. Sequences were aligned (Mafft, L-INS-I, “leave gappy regions”) and 392 informative residues were selected (Gblocks, relaxed parameters) for tree inference (PhyML, GTR model, SH-aLRT with 1000 replicates). Sequences from non-Helotiales Leotiomycetes were used for rooting (grey triangle). When available, the fungal lifestyle is indicated (as in Bruyant *et al*. 2024). Plant pathogens are described as biotrophs (i.e. they colonize living plant tissue and obtain nutrients from living host cells) or necrotrophs (i.e. they colonize living plant tissue and obtain nutrients from dead tissues). Symbiotic Helotiales are marked as either forming ericoid mycorrhiza or ectomycorrhiza, and root endophytes are characterized as endophytes with no known effect on plants or as beneficial root endophytes. Purple NT symbols indicate strains shown to transfer nutrients to their hosts, and mycorrhizal-like strains are highlighted in turquoise. The relative abundance of isolates in the collection or in the roots of plants by metabarcoding is indicating by heatmaps. Blue dots on branches indicate bootstrap > 0.8. The branch lengths are not proportional to the amount of evolutionary change or time.

In the Hyaloscyphoid clade, two isolates were found to cluster with plant-beneficial strains, such as ericoid mycorrhizal strains *Meliniomyces bicolor* E, and ectomycorrhizal strain *Cadophora finlandica* isolate PRF15. However, none of our isolates were closely related to the mycorrhizal-like species *Pezoloma ericae* associated with the liverwort *Cephalozia bicuspidata*.

The Mollisioid clade included nine isolates. This clade featured isolates closely related to mycorrhizal-like *Ascomycota* sp. Da MaP I9 *Tapesia* sp., which transfers nitrogen to *Deschampsia antarctica* (Poaceae), and *Helotiales* sp. F229, a *Cadophora*-related strain that facilitates phosphorus transfer to non-mycorrhizal *Arabis alpina*. Additionally, we identified isolates close to plant growth-promoting strains from the *Lepdodophora/Leptodontium* group and *Phialocephala* group, but did not recover isolates closely related to pathogenic species from e.g. the *Diplocarpon* or *Marssonina* genera.

The Pezizzeloid clade encompassed most isolates (16 out of 30). It included two *Pezicula* isolates and two *Microscypha* isolates, both closely related to plant-beneficial endophytes. Additionally, four isolates from the *Lachnum* genus were identified. This genus contains dark septate endophytes and endophytes with growth-promoting effects on *Vaccinium* species. Interestingly, we also found a *Titaea maxilliformis* isolate related to the lichen-associated strain *Pezizella epithallina*. Despite the high prevalence of this lineage in plant roots (ITS2 metabarcoding), only one isolate was recovered. The other isolates from this clade were close to strains with undefined lifestyles or to saprotrophs. We can also note the absence of isolates related to ericoid mycorrhizal fungi, such as *Oidiodendron maius* or *Gamarada debralockiae*, which was expected as these strains are exclusively described as ericoid-forming fungi.

Unexpectedly, the Discinelloid clade included two isolates that are not closely related to known root endophytes but are described as saprotrophs, which were also not detected in the metabarcoding analyses. Additionally, no isolates from the Sclerotinoid clade were recovered. This clade encompasses pathogenic or lichen-associated species, and their absence aligns with our expectation given the healthy condition of the plants and low prevalence of these taxa in metabarcoding analyses.

In summary, our culture method captured a significant fraction of the Helotiales diversity, with a notable majority of them belonging to the Pezizzeloid clade, where mycorrhizal-like fungi have not yet been described. Additionally, we successfully retrieved some isolates close to mycorrhizal-like lineages from the Mollisioid clade.

### P solubilization potential of Helotiales and other fungal isolates

Finally, we assessed the phosphate solubilization potential of selected isolates in vitro, hypothesizing that this capacity might be prevalent among root-endophytic Helotiales due to their potential role in enhancing P nutrition for non-mycorrhizal (Non-AM) plants. We screened 69 Helotiales isolates and 73 non-Helotiales isolates from our collection for phosphorus solubilization activity using PVK medium (Table S2). Surprisingly, only 16 isolates (11.3%) exhibited phosphate solubilization *in vitro*. These included strains from the Hypocreales (8 *Fusarium* sp. or closely related strains), Helotiales (6 isolates, among which 4 belonged to the *Pezicula* genus, each from a different plant species, 1 was closely related to *Cadophora* strains, and 1 was a *Lachnum* sp.), and Pleosporales (1 *Alternaria* sp. and 1 *Montagnula* sp.). The *Pezicula* genus (Helotiales) is particularly noteworthy, as half of the tested isolates could solubilize phosphate, making them especially promising as potential enhancers of plant nutrition. In summary, while phosphate solubilization potential was demonstrated in a minority of fungal isolates, even Helotiales ones, the presence of this capability in several Helotiales *Pezicula* strains underscores their potential significance for enhancing plant P nutrition.

## Discussion

As root endophytes, Helotiales fungi might play a critical role in supporting plant nutrition in Arctic-alpine regions, but their exact ecological function remains unclear. While these fungi are hypothesized to be particularly crucial for the nutrition of Non-AM plants in such environments (Almario *et al*., 2022; Bruyant *et al*., 2024). Their ecology remains largely unknown, as relatively, few strains have been successfully isolated and studied. This study sought to assess the efficacy of a tailored culture-based method for exploring the taxonomic diversity of root-endophytic Helotiales within the roots of alpine Non-AM plants, with a specific focus on lineages with established plant-beneficial potential.

A major challenge in plant microbiota research is the curation of microbial culture collections capturing the diversity in natural plant microbiota (Liu *et al*., 2022). Here, from the 3,920 surface-sterilized root fragments plated and 1,733 fungal isolates screened, only 602 distinct isolates were obtained, representing approximately one-third (31.1%) of the fungal root microbiota of these plants. This mirrors results from previous studies in other plants (Glynou *et al*., 2016, 2018; Durán *et al*., 2018; Bonito *et al*., 2016), and suggests that despite great sampling efforts, capturing more than a third of root-associated fungal taxa as pure cultures remains extremely challenging.

In comparison, Helotiales taxa were much better captured in our collection with approximately 42.8% of Helotiales taxa represented (Fig. 2). In comparison with studies by Glynou *et al*. (2016) and Durán *et al*. (2018), our approach yielded a higher proportion of these fungi in collection (53.0% vs 18.2% and 1.2%, respectively). The higher proportion of Helotiales retrieved in our collection might be due to the use of diverse plant species. Indeed, we obtained most Helotiales isolates from Cyperaceae plants, which also harbored higher Helotiales diversity. While Brassicaceae have been noted as a hotspot for Helotiales diversity (Maciá-Vicente *et al*., 2020), our study suggests Cyperaceae are too. It is also possible that roots of alpine plants, growing in nutrient poor environments are in general enriched in Helotiales, as several metabarcoding analyses found that root Helotiales are particularly abundant in these environments (Toju *et al*., 2016; Almario *et al*., 2017; Jumpponen *et al*., 2017; Botnen *et al*., 2020). Contrary to expectations, the three culture media used for isolation recovered a similar fungal diversity, even within Helotiales. Media with reduced complexity (such as MM) are often used to support the growth of slow-growing species (dos Reis *et al*., 2022; Hagh-Doust *et al*., 2022) rather than complex ones such as Potato Dextrose Agar (PDA), while Hagem’s medium (HgM; without benlate) was originally developed to recover ectomycorrhizal or ericoid mycorrhizal fungi (Walker *et al*., 2011; Rämä and Quandt, 2021; Hagh-Doust *et al*., 2022), including Helotiales. Interestingly, only the nutrient-poor synthetic medium (MM) allowed the recovery of Helotiales fungi across all screened plants. This observation may be attributed to the characteristics of cold alpine environments, which are typically low in nutrients (Vincent *et al*., 2014) and are known to host many slow-growing, oligotrophic fungi (Godinho *et al*., 2015). Helotiales fungi, in particular, encompass many water fungi and are abundant in these nutrient-poor settings (Toju *et al*., 2016; Jumpponen *et al*., 2017; Botnen *et al*., 2020; Almario *et al*., 2022) suggesting they are adapted to low nutrient environments.

Our approach showed high phylogenetic diversity among cultured root-endophytic Helotiales, capturing five of the six known clades, as found by Glynou *et al*. (2016) in European Brassicaceae. This is particularly notable given that Glynou *et al*. (2016) studied 52 plant populations, whereas our study focused only on six populations. In comparison, the study of Walker *et al*. (2011) targeting Arctic Ericaceae, isolated several Helotiales but only from three clades (Mollisioid, Hyaloscyphoid, and Pezizzeloid), while studies from non-alpine Brassicaceae have cultured only specific genera, such as *Leptodontidium* (Mollisioid clade) (Durán *et al*., 2018), *Phialophora* (Mollisioid clade) and *Tetracladium* (Pezizzeloid clade) (Keim *et al*., 2014). This comparison underscores the added value of our culture approach, which effectively captured a broad spectrum of root-associated Helotiales lineages.

Most cultured root-endophytic Helotiales lineages (28 of 30 representative isolates) belonged to clades known to include several plant-beneficial lineages. Within the Mollisioid clade we recovered two isolates from *Carex* roots that are closely related to the mycorrhizal-like *Tapesia* strain I9 (*Ascomycota* sp. strain Da MaP I9), known to transfer N to its antarctic host, *Deschampsia antarctica* (Hill *et al*., 2019) (monocot, Poaceae). These Helotiales could similarly sustain N nutrition in Cyperaceae (monocots) growing in arctic-alpine environments. Within this same clade, we recovered as many as 59 isolates closely related to the mycorrhizal-like strain Helotiales sp. F229 from all plant species except *M. verna* (Caryophyllaceae of Galibier). Helotiales sp. F229 was shown to transfer P to *A. alpina* (Almario *et al*., 2017), suggesting these lineage might promote P nutrition in other alpine non-AM plants.

In the Hyaloscyphoid clade, we found 6 isolates close to the ericoid mycorrhizal *Hyaloscypha bicolor* E, known to transfer N to its Ericaceae hosts (Grelet *et al*., 2009). Although it is not known if these fungi can transfer N to non-Ericaceae plants, we could imagine they may act as mycorrhizal-like partners promoting N nutrition in non-AM plants. We did not find however, isolates closely related to mycorrhizal-like fungus *Pezeloma ericae*, suggesting that this lineage might either challenging to cultivate or absent from our sampling conditions.

In the Pezizzeloid clade, we recovered one isolate from the highly abundant Helotiales taxon *Titaea maxilliformis* (syn. *Tetracladium maxilliforme*; Pezizzeloid clade). *Tetracladium* species are mostly known as water fungi but have been recently linked to increased yields in the non-AM crop *Brassica napus* (Lazar *et al*., 2022; Li *et al*., 2023). Plant-beneficial functions in these fungi are however poorly described. Interestingly, as many as 78 *Pezicula* were isolated in our collection, with several isolates solubilising phosphate *in vitro*, as shown elsewhere (Luo *et al*., 2023; Dasila *et al*., 2024). This suggests this lineage might harbour undescribed fungi with plant beneficial activities involving improved plant P-nutrition.

This study presents an effective method for isolating and cultivating beneficial root-endophytic Helotiales, demonstrating that alpine non-mycorrhizal plants are a valuable source of these fungi. Our collection of fungal strains provides crucial biological material for further investigating the mechanisms underlying Helotiales-plant interactions.

## Acknowledgements

This work was supported by Lautaret Garden – UAR 3370 (Univ. Grenoble Alpes, CNRS, 38000 Grenoble, France), a member of AnaEE France (ANR-11-INBS-0001 AnaEE-Services, Investissements d’Avenir frame) and of the eLTER European network project (Zone atelier Alpes, CNRS, UGA 38000 Grenoble France). We thank specially Maxime Rome (Lautaret Garden). This work was performed using the computing facilities of the CC LBBE/PRABI-AMSB and the IBIO platform (LEM UMR5557).

## Data Availability

The authors confirm that all data underlying the findings are fully available without restriction. All sequencing data (i.e. generated either by the Sanger sequencing or by Illumina Miseq technology) are deposited at the NCBI database under the following accession numbers: PQ286942-PQ286971 for complete ITS Sanger sequences and PQ323580 - PQ324178 for the ITS2 Sanger sequences and PRJNA1141433 for the fastq-files generated by the Illumina (Miseq).

## Author information

### Contributions

J.A., P.B. and Y.M.L. initiated the research, conceived and designed the experiments. J.A., J.D and P.B. collected wild-growing plants. P.B., J.D. and L.V. created the fungal culture collection. P.B., J.A. and Y.M.L. wrote the manuscript with help from all other co-authors.

## Ethics declarations

### Ethics approval and consent to participate

Not applicable.

### Consent for publication

Not applicable.

## Supplementary tables

Supplementary file: File Excel_Supp_Tables_.xlsx

TABLE S1. Soil properties of the sampling sites

TABLE S2. List of isolates obtained.

TABLE S3. Number of root segments giving fungal isolates and numbers of isolates sequences.

